# Mechanistic basis for non-exponential bacterial growth

**DOI:** 10.1101/2025.03.29.646116

**Authors:** Arianna Cylke, Shiladitya Banerjee

## Abstract

Bacterial populations typically exhibit exponential growth under resource-rich conditions, yet individual cells often deviate from this pattern. Recent work has shown that the elongation rates of *Escherichia coli* and *Caulobacter crescentus* increase throughout the cell cycle (super-exponential growth), while *Bacillus subtilis* displays a mid-cycle minimum (convex growth), and *Mycobacterium tuberculosis* grows linearly. Here, we develop a single-cell model linking gene expression, proteome allocation, and mass growth to explain these diverse growth trajectories. By calibrating model parameters with experimental data, we show that DNA-proportional mRNA transcription produces near-exponential growth, whereas deviations from this proportionality yield the observed non-exponential growth patterns. Analysis of gene expression perturbations reveals that ribosome expression primarily controls dry mass growth rate, whereas envelope expression more strongly affects cell elongation rate. Fitting our model to single-cell experimental data reproduces convex, super-exponential, and linear modes of growth, demonstrating how envelope and ribosome expression schedules drive cell-cycle-specific behaviors. These findings provide a mechanistic basis for non-exponential single-cell growth and offer insights into how bacterial cells dynamically regulate elongation rates within each generation.

## I. INTRODUCTION

Early studies in bacterial growth physiology established that bacterial populations grow exponentially under nutrient-rich conditions where resources are not limiting [1]. Recent advancements in microfluidics and single-cell imaging technologies have enabled the collection of extensive, high-resolution single-cell datasets for bacterial growth [2–8]. These datasets have fueled the development of models addressing single-cell division control and size homeostasis [9–14], cell size control [13, 15–17], shape maintenance [12, 18–20], and adaptation to environmental changes [21–25]. Many of these models either assume exponential growth or overlook intra-generational dynamics. However, detailed analysis of high-resolution time-series data has revealed that elongation rates are not constant throughout the bacterial cell cycle. For instance, *Escherichia coli* [26, 27] *and Caulobacter crescentus* [27, 28] exhibit super-exponential growth, characterized by increasing elongation rates over the cell cycle. In contrast, *Bacillus subtilis* displays a convex elongation rate pattern, with a minimum rate near the mid-cell cycle [29], while *Mycobacterium tuberculosis* grows linearly [30]. To the best of our knowledge, no bacterial species has been reported to exhibit truly exponential elongation at the single-cell level. These observations raise the fundamental question of what drives these time-dependent growth patterns.

Cell growth is fundamentally driven by the translation of new proteins by ribosomes, with a variety of factors such as animo acid precursor and ribosome abundances influencing the rate at which overall protein translation occurs. Resource allocation models for the cellular proteome have been shown to be widely applicable for understanding bacterial growth from steady-state [17, 31–34] to dynamic environments [25, 35–38]. According to the central dogma of molecular biology, genes are transcribed into mRNA, which is then translated by ribosomes to produce the protein mass of the cell [39]. For balanced exponential growth, the relative expression of genes determines the composition of the cell’s proteome in a given environment [40], with constant gene expression and stable concentrations of mRNA and proteins [41]. Intuitively, non-exponential growth must involve deviations from these constant concentrations over the course of the cell cycle. Recent studies have revealed that transcription is linked to DNA replication, with transient increases in transcription occurring during the replication of specific genes [42, 43]. Additionally, transcription rate is often assumed to be proportional to the DNA content of the cell [40, 43]. Given that the duration of DNA replication does not always align with the cell cycle time [8], deviations from balanced exponential growth may not be surprising.

In this work, we examine how cell-cycle-dependent gene expression influences single-cell growth by combining proteome allocation theory and mRNA dynamics into a unified model. Building on existing models and guided by experimental data, we formulate a framework that explicitly includes two key sectors—ribosomes and the cell envelope—treating nutrient availability and mRNA transcription rates as the primary free parameters. Initially, we find that transcription scaled to DNA content yields near-exponential growth, even with DNA concentration fluctuations throughout the cell cycle. However, this prediction does not match the non-exponential growth patterns observed in many bacterial species. Consequently, we introduce additional time-dependent transcription dynamics to better capture experimental findings. By systematically varying transcription rates, we quantify how these perturbations shape cell growth, both in general and within the context of cell cycle data. We further pinpoint the timescale limitations that govern how gene expression and downstream processes affect overall growth. Using these insights, we explain the mechanistic origins of non-monotonic elongation rate in *B. subtilis*, super-exponential growth in *E. coli*, and linear growth in *M. tuberculosis*, underscoring how trade-offs between ribosome and envelope synthesis vary over the cell cycle. Together, these findings provide a mechanistic understanding of the regulatory principles shaping bacterial growth dynamics at the single-cell level.

## II. NON-EXPONENTIAL ELONGATION MODES OF SINGLE BACTERIAL CELLS

While bacterial population growth is often quantified by mass or optical density, single-cell growth data are collected via microscopy and image analysis. Consequently, single-cell growth metrics typically focus on morphological parameters, such as cell length and width. For the rod-like bacterial cells examined in this study, cell width remains approximately constant throughout the cell cycle; changes in cell length directly reflect growth. Hence, we characterize single-cell growth by the elongation rate, *κ*_*L*_ = *L*^−1^*dL/dt*, which is constant for purely exponential growth. In practice, however, experimental time-lapse data do not show purely exponential growth in cell length trajectories.

Figure 1a shows qualitative examples of super-exponential (*E. coli*), convex (*B. subtilis*), and linear (*M. tuberculosis*) elongation rates, while Fig.1b displays representative datasets from these species [4, 8, 12, 30]. To facilitate cross-species comparisons—given their distinct growth timescales—we plot the fractional deviation from the mean elongation rate, 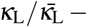 1. Here, super-exponential growth appears concave and asymptotically approaches exponential behavior [27], whereas linear growth is convex. Notably, in *B. subtilis*, the convex growth pattern begins and ends each generation at approximately the same elongation rate, whereas super-exponential and linear growth modes are strictly monotonic, resetting to an initial rate after division. It is important to recognize that non-exponential growth at the single-cell level does not necessarily imply non-exponential population growth; as long as the cell cycle duration remains consistent, the overall population still grows exponentially.

**FIG. 1.**
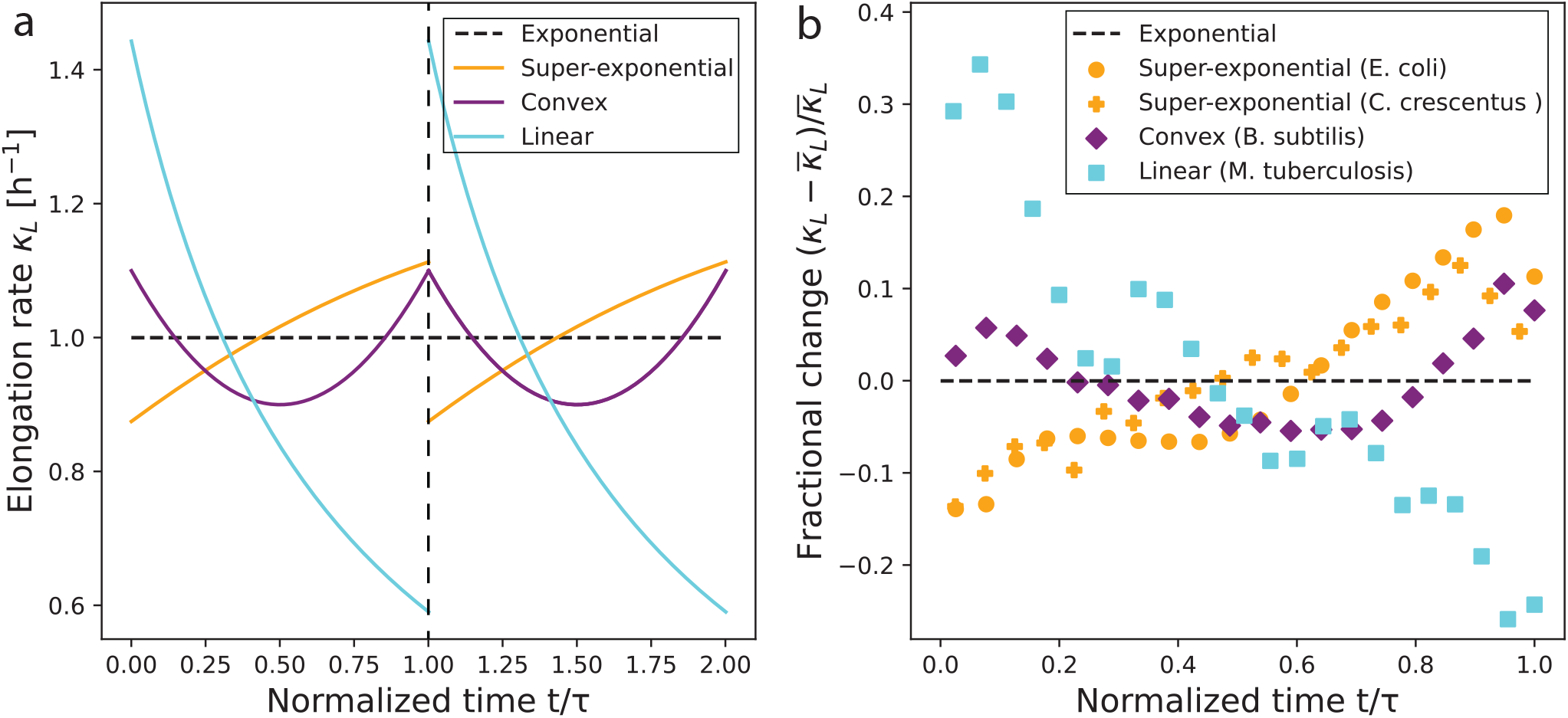
Non-exponential elongation modes in bacterial cells. (a) Qualitative growth trajectories illustrating how various modes of bacterial growth deviate from purely exponential growth. The elongation rate *κ*_*L*_ = *L*^−1^*dL/dt* is plotted against normalized time *t/τ* for two cell cycles, where *τ* is cell cycle duration. (b) Ensemble-averaged elongation rate data for four representative bacterial cells of the growth modes illustrated in (a), plotted in normalized time. Individual cell elongation rate trajectories were extracted from length data via a trapezoidal derivative approximation and then averaged in bins of normalized time. For ease of comparison, we plot the fractional change in growth rate (where 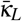 denotes cell cycle average). *E. coli* K12 NCM3722 cells are grown in supplemented Glucose media at 37°C [4], *C. crescentus* cells are grown in PYE at 34°C [12], *B. subtilis* BSB1 cells are grown in Glycerol at 37°C [8], and *M. tuberculosis* CDC1551 cells are grown in supplemented 7H9 growth medium at 37°C [30].

Despite these observations, our mechanistic understanding of non-exponential growth remains limited. Existing models of super-exponential and linear growth largely focus on cell-envelope processes, suggesting that *E. coli* elongates exponentially in only a subsection of the cell envelope [27], whereas *M. tuberculosis* grows exclusively at its poles [30]. For *B. subtilis*, the location of the mid-cycle minimum in elongation rate has been linked to cell size control [8]. However, these morphological perspectives do not fully explain the underlying physiological mechanisms driving non-exponential growth. In particular, what do these patterns imply about gene expression and proteome changes within a single cell cycle? In subsequent sections, we place these distinct growth modes in the context of the central dogma of molecular biology, aiming to understand how non-exponential single-cell growth arises from temporal patterns of gene expression and proteome allocation.

## III. MECHANISTIC MODEL FOR SINGLE-CELL GROWTH CONTROL

To explain the mechanistic basis for exponential and non-exponential bacterial growth at the single-cell level, we develop a theoretical framework that links gene expression to proteome allocation and nutrient uptake, ultimately driving single-cell growth (Fig. 2). We begin by adapting the well-established proteome allocation theory, which partitions the cellular proteome into ribosomal, envelope, and other protein sectors. We then couple this allocation to a dynamical description of mRNA transcription, ribosome binding, and protein synthesis. Next, we incorporate the role of DNA replication in modulating transcription rates and thus gene dosage. Finally, we outline how these biochemical processes translate into changes in cell morphology and growth, enabling direct comparison with experimental data. Together, these components provide a comprehensive mechanistic model of how time-dependent gene expression shapes bacterial growth at the single-cell level.

**FIG. 2.**
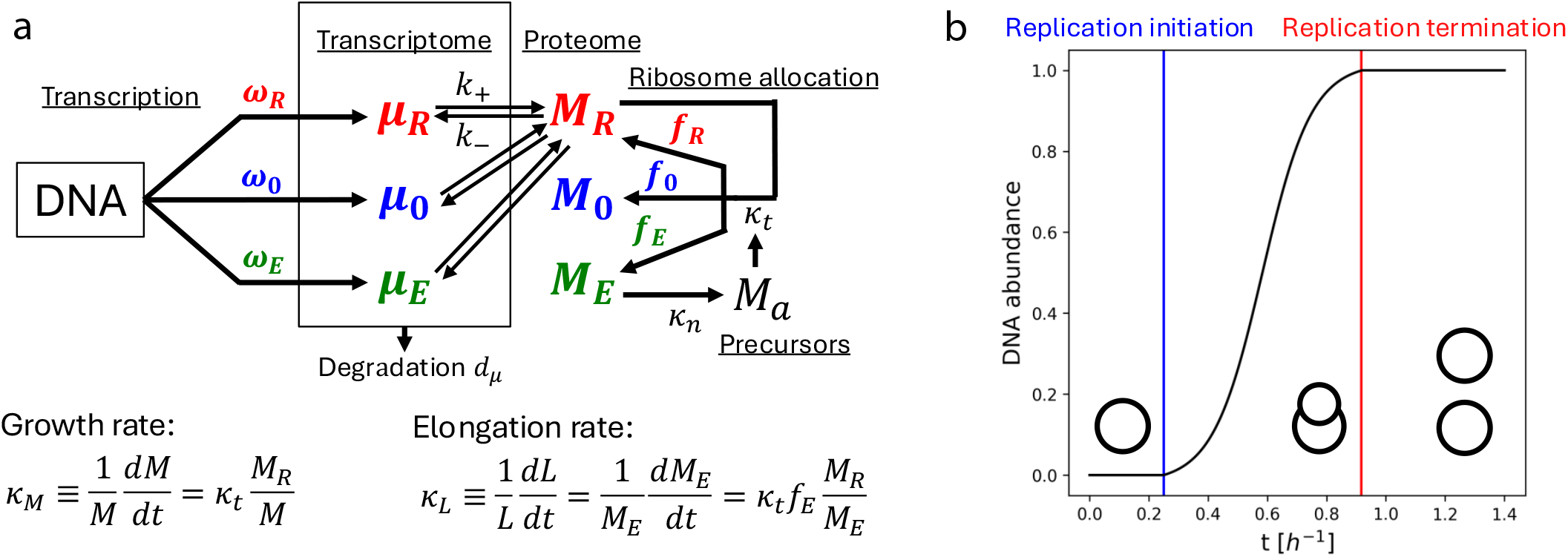
Overview of the single-cell growth model. (a) Schematic illustrating how DNA abundance determines mRNA transcription rates, which then set mRNA levels for each proteome sector. The mRNA binds to ribomsomes to determine the allocation fractions for each sector ribosomal, envelope, or “other”) and degrades over time when unbound. Ribosomes grow the dry mass of the cell through translation of proteins in proportions set by the allocation fractions. The cell envelope mass sets the surface area and therefore the rate at which nutrients are imported, the abundance of which sets the transcription rate. (b) A sample DNA-replication logistic function describing DNA abundance in absolute time, as defined in Eq. (12). Blue and red vertical lines mark replication initiation and termination, respectively, and the black circles depict the bacterial chromosome at different replication stages.

### A. Proteome allocation

We begin by adopting a well-established proteome allocation framework for bacterial growth [31, 32]. In such models, cell dry mass *M* increases through the translation of new proteins by ribosomes. If the translation rate is *κ*_*t*_, then

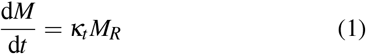

where *M*_*R*_ is the abundance of ribosomes. Cell growth rate is defined as the relative growth rate of total cell mass:

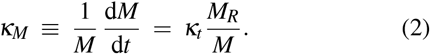

In our model, we divide the proteome into three main sectors: (i) ribosomal proteins of mass *M*_*R*_, the cell envelope (and periplasm) of mass *M*_*E*_, and all other proteins of mass *M*_0_ ≡*M* −*M*_*R*_ −*M*_*E*_. We define an allocation fraction *f*_*q*_ as the fraction of actively translating ribosomes devoted to synthesizing sector *q*. Accordingly,

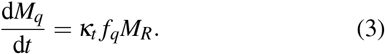

Ribosomes not actively translating any sector are accounted for by a free ribosome fraction *f*_*F*_. Thus, *f*_*R*_ + *f*_*E*_ + *f*_0_ + *f*_*F*_ = 1 spans the entire proteome’s translational allocation.

To track the changing distribution of mass between sectors, we introduce the mass fraction *φ*_*q*_ = *M*_*q*_*/M*. Combining the definition of *φ*_*q*_ with Eq. (3), we arrive at

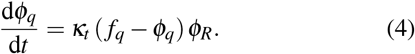

Under steady-state conditions, constant allocations imply *f*_*q*_ = *φ*_*q*_. However, for time-varying *f*_*q*_(*t*), the instantaneous *φ*_*q*_ lags behind *f*_*q*_, relaxing toward it at a rate set by *κ*_*t*_*φ*_*R*_. As long as these allocation changes are cyclic and repeat each generation, the inter-generational average mass fractions 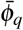 remain consistent — even if fluctuations arise within the cell cycle.

Bacterial cells can be limited by nutrient uptake, translation capacity, or a combination of the two. Cells grow optimally when they balance both of these limitations to maximize overall growth rather than translation speed or precursor abundance [32]. Our model incorporates these limitations by introducing an amino acid precursor pool of mass *M*_*a*_. Flux balance implies:

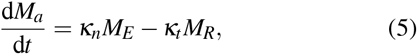

where *κ*_*n*_ denotes the nutrient uptake rate. We assume that nutrient uptake is proportional to the cell envelope mass, consistent with uniform nutrient transport across the cell surface [44, 45].

The translation rate *κ*_*t*_ decreases when the amino acid mass fraction *φ*_*a*_ = *M*_*a*_*/M* is low, reflecting precursor scarcity. We capture this using

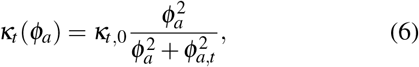

where *κ*_*t*,0_ is the maximum translation rate, and *φ*_*a,t*_ sets the threshold at which precursor limitation attenuates translation. Similarly, nutrient uptake undergoes feedback inhibition at high *φ*_*a*_:

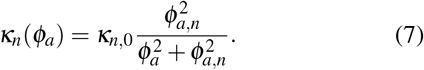

with *φ*_*a,n*_ determining the threshold for uptake feedback and *κ*_*n*,0_ the maximal uptake rate. Parameter values estimated for *E. coli* inclues: *φ*_*a,t*_ = 10^−4^, *φ*_*a,n*_ = 10^−3^, and 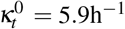 [32].

Although we primarily focus on cells in nutrient-rich conditions, explicitly including precursors in the model highlights the trade-off between ribosomes and the envelope. Allocating more resources to ribosomes increases the cell’s overall translation capacity, while reducing envelope allocation limits nutrient uptake. Thus, our framework captures both translation-driven and nutrient-driven constraints in single-cell growth.

### B. Gene expression dynamics

Allocation of proteome resources within the cell is governed by gene expression. In a minimal model, DNA is transcribed into mRNA, which then provides instructions for ribosomes to synthesize proteins. To connect gene expression to dynamic proteome allocation, we adapt the framework from Ref. [34], defining *µ*_*q*_ as the mRNA abundance for each proteome sector *q* ∈ {*R, E*, 0}. The corresponding time-evolution equation is:

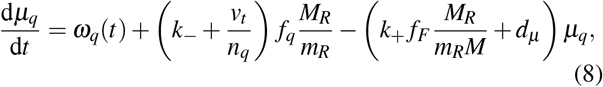

where *ω*_*q*_(*t*) is the time-dependent mRNA transcription rate, *k*_+_ (*k*_−_) is the rate of mRNA-ribosome binding (unbinding), *v*_*t*_ is the rate at which a single ribosome translates amino acids, *n*_*q*_ is the average number of amino acids per protein in the sector, *m*_*R*_ is the mass of a single ribosome, and *d*_*µ*_ is the mRNA degradation rate. Note that the term *v*_*t*_*/n*_*q*_ corresponds to ribosome unbinding upon completion of protein synthesis. The equation governing the time-evolution of mRNA-ribosome complex is given by:

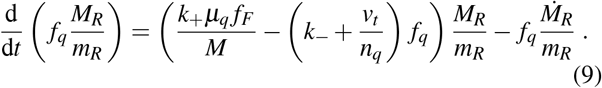

Using Eq. (3) to substitute for 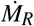, we derive the time dependence of *f*_*q*_:

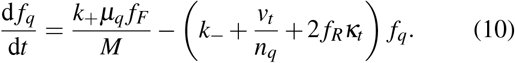

Since ∑_*q*_ *f*_*q*_ = 1, we also have,

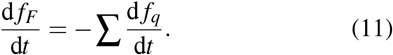

Most of these parameters are known for *E. coli*: *d*_*µ*_ = 6h^−1^ [46], *k* = 60h^−1^ and *k*_+_ = 1.1 10^5^MDa/h [34]. Ribosomes contain *n*_*R*_ ≈ 7459 amino acids [47], so *v*_*t*_ = *n*_*R*_ *κ*_*t*_, and each ribosome has a mass *m*_*R*_ ≈ 2.7 MDa [48]. Typical envelope or general proteome proteins have ~ 287–294 amino acids in *E. coli* and *B. subtilis* [49]. While *n*_*R*_ describes a complete ribosome, ribosome-affiliated proteins are translated individually, and thus we approximate *n*_*E*_ = *n*_0_ = *n*_*R*_ in Eq. (10). Consequently, the only remaining free parameters are the transcription rates *ω*_*q*_(*t*), the maximal nutrient uptake *κ*_*n*,0_, and the initial conditions used to integrate the system.

Figure 2a summarizes this framework in a schematic: DNA abundance sets the mRNA transcription rates *ω*_*q*_(*t*), forming ribosome-mRNA complexes for each proteome sector (ribosomal, envelope, and “other”). The cell envelope then modulates nutrient uptake, which in turn affects the cell’s amino acid pool and, consequently, translation. Figure 2b shows a sample logistic function for DNA abundance, with blue and red vertical lines marking initiation and termination of DNA replication that shape these transcription dynamics. Overall, this model links gene expression (via mRNA transcription) to proteome allocation, enabling us to study how changing transcription rates throughout the cell cycle impact bacterial growth dynamics.

### C. DNA replication dynamics

Experiments indicate that transcription is roughly proportional to gene dosage, i.e., to the cell’s DNA content [40, 43]. Bacterial cells maintain a tightly controlled cell volume per replication origin, ensuring that DNA replication initiates at a specific cell size regardless of nutrient availability. For *E. coli*, this critical volume per origin is about 0.28 *µ*m^3^, while for *B. subtilis* it is approximately 0.60 *µ*m^3^ [8].

Once replication begins, the duration of the *C* period (time interval between the initiation and termination of chromosome replication), *τ*_*C*_, can be relatively growth-rate-independent under nutrient-rich conditions (i.e., for *κ* ⪆ 0.7 h^−1^). In *E. coli, τ*_*C*_ has been measured at 39 min [8], whereas *B. subtilis* exhibits a similar baseline of 40 min (though it shows a weak linear dependence on growth rate, fitted by 1.02 h^−1^ − 0.26 *κ* [8]).

Using these two parameters—the initiation time and the replication duration—we model the abundance of a single chromosome by a logistic function:

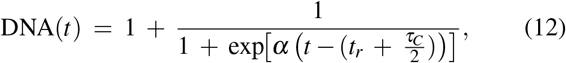

where *t*_*r*_ is the time at which the initiation criterion (critical cell volume) is met, and 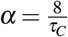 ensures that ~ 98% of replication completes within the interval [*t*_*r*_, *t*_*r*_ + *τ*_*C*_]. Consequently, the instantaneous transcription rate for mRNA in sector *q* can be written as

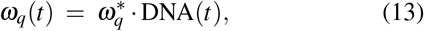

where 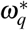 is the transcription rate per chromosome. We term this *DNA-proportional transcription* model. Figure 2b shows a sample logistic curve illustrating this replication-driven modulation of *ω*_*q*_(*t*) during one cell cycle.

### D. Cell morphology and growth

To compare our model predictions with microscopy-based measurements, we approximate each bacterial cell as a cylindrical rod of length *L* and width *w*, with hemispherical end caps. The cell volume is given by

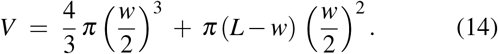

We can convert between cell morphology and mass using experimental measurements for the average dry mass density *ρ*. For instance, *E. coli* has *ρ*≈ 0.30 gml^−1^ ≈1.7× 10^5^ MDa *µ*m^−3^ [20], while *B. subtilis* has *ρ* ≈0.32 gml^−1^≈ 1.9 × 10^5^ MDa *µ*m^−3^ [50].

In rod-like bacteria, the cell envelope mass *M*_*E*_ is proportional to the total surface area of the cylinder and hemispherical end caps, *S* = *π w L*, which we write as *M*_*E*_ = *d ρ*_*E*_ *π wL*, where *ρ*_*E*_ is the envelope density and *d* its thickness. Consequently, the increase in *M*_*E*_ reflects the increase in *L*, so the cell’s elongation rate can be expressed as

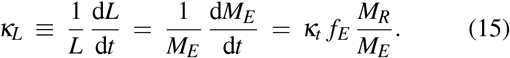

This definition allows direct comparison to experimental measurements of cell length. By contrast, the growth rate of total cell mass is

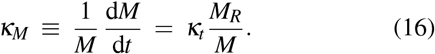

Hence, *κ*_*L*_ quantifies cell size elongation while *κ*_*M*_ quantifies how total cellular biomass increases over time.

## IV. MODEL IMPLEMENTATION: EMERGENCE OF EXPONENTIAL GROWTH

We now describe how our single-cell growth model is numerically implemented, including the implementation of cell division rule and intergenerational dynamics of cellular proteome and transcriptome. Using this model, we show that transcription rates scaling with cellular DNA content naturally lead to near-exponential growth, even when DNA replication is not strictly synchronized with cell mass.

To simulate how single cells evolve their size, growth rate, and proteome composition across generations, we explicitly incorporate cell division in our simulations. At the beginning of the first simulated cell cycle, we specify the initial values for each sector’s mRNA abundance *µ*_*q*_, total mass *M*_*q*_, allocation fractions *f*_*q*_, and mass fractions *φ*_*q*_. These quantities then converge to their homeostatic values over multiple generations, consistent with the homeostasis in cell size and growth rate. In particular, we implement an “adder” rule for cell size regulation and division control, supported by experimental data showing that rod-shaped bacteria add a constant length (and therefore a constant envelope mass *M*_*E*_) between divisions, regardless of the birth length [4, 8, 10]. Once the cell meets its division criterion, we implement symmetric division by halving each extrinsic quantity (*µ*_*q*_, *M*_*q*_, and *L*) in daughter cells. The final values of *f*_*q*_ and *φ*_*q*_ at division become the initial values for the daughter cells in the next generation.

Next, we detail how the model equations are integrated and how DNA replication is coupled to transcription. At each timestep during the cell cycle, we dynamically update:

- The DNA content, using Eq. (12), whenever the cell meets the initiation mass threshold. For cells undergoing multifork replication, the process is identical with summed contributions from each chromosome.
- The sector-specific mRNA abundances *µ*_*q*_(*t*) via Eq. (8), where 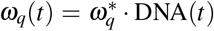sets the instantaneous transcription rate;
- The proteome allocation fractions *f*_*q*_(*t*) using Eqs. (10) and (11);
- The sector masses *M*_*q*_(*t*) using Eqs. (1) and (5);
- The mass fractions *φ*_*q*_(*t*) = *M*_*q*_(*t*)*/M*(*t*) using Eq. (4).

If multifork replication has not completed by the time of cell division, we allow the unfinished replication forks to continue in the next generation, assuming equal partitioning of chromosomes upon division. After cell division, if all state variables (*M*_*q*_, *µ*_*q*_, *f*_*q*_, *φ*_*q*_) match their values at the start of the previous cycle within a small tolerance (e.g., 1%), we consider the system to have reached a steady-state cycle.

In practice, we tune the three per-chromosome transcription rates 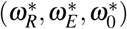 and the maximal nutrient uptake rate *κ*_*n*,0_ so that the steady-state cell cycle time *τ*, the average envelope fraction 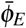, and the mean free ribosome fraction 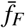 match experimental data. For *E. coli*, 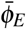 typically falls between 0.058 and 0.152 depending on growth rate [51], while *B. subtilis* in rich media has 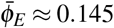 regardless of growth rate [50]. The free ribosome mass fraction 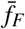 is approximately 0.05 in exponentially growing *E. coli* [31], a value we also adopt for *B. subtilis*.

An example simulation of the DNA-proportional transcription model is shown in Fig. 3 for *B. subtilis* growing in glycerol-rich media with 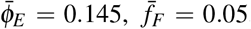, and *τ* = 0.41 h. Under these conditions, 4–8 replication origins are present at any given time. As the DNA content increases, the transcription rates *ω*_*q*_(*t*) and the corresponding mRNA abundances *µ*_*q*_(*t*) rise throughout the cell cycle (Fig. 3a). However, the allocation fractions *f*_*q*_(*t*) and mass fractions *φ*_*q*_(*t*) remain nearly constant (Fig. 3a). Consequently, both the mass growth rate *κ*_*M*_ and elongation rate *κ*_*L*_ exhibit only small amplitude fluctuations (less than 0.3% from their means), leading to near-exponential growth over each cell cycle (Fig. 3b).

**FIG. 3.**
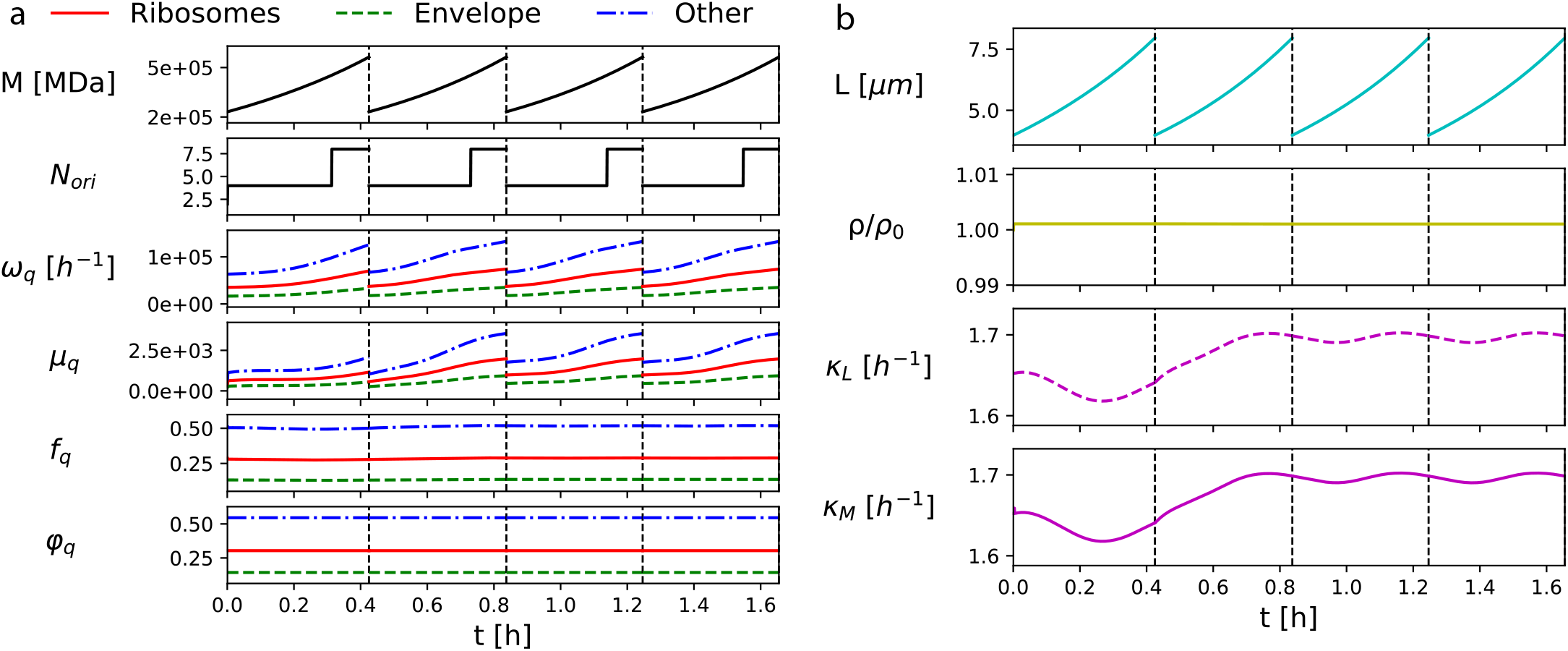
Representative simulation of the DNA-proprtional transcription model, showing the evolution of cell physiological variables. (a) Trajectories of physiological variables evolved in the model (top to bottom): cell mass *M*, the number of replication origins *N*_*ori*_, transcription rates *ω*_*q*_, mRNA abundance *µ*_*q*_, proteomme allocation fraction *f*_*q*_ and mass fraction *φ*_*q*_. Step increase in replication origins per cell denotes replication initiation. The dashed vertical line denotes a cell division event. The parameters used are *κ*_*n*,0_ = 25h^−1^, *ω*_*R*_ = 1.77 · 10^4^h^−1^, *ω*^*E*^ = 8.45 ·10^3^h^−1^, and *ω*_0_ = 3.18 ·10^4^h^−1^ with a steady-state tolerance window of 1%. (b) Model output showing the trajectory of (top to bottom) cell length *L*, normalized cell density *ρ/ρ*_0_, elongation rate *κ*_*L*_, and mass growth rate *κ*_*M*_ for the simulation corresponding to data in panel (a). Density is relative to the initialized value *ρ* = 1.9 ·10^5^MDa*/µ*m^3^ [50]. Elongation and growth rates are calculated according to Eq. (15) and Eq. (16), respectively.

Formally, purely exponential single-cell growth would require that the DNA content and, thus, transcription rates be strictly proportional to cell mass: DNA(*t*) = DNA(0) *M*(*t*)*/M*(0). Any departure from this proportionality introduces small oscillations in the instantaneous growth rate about the equilibrium value. Nevertheless, our results show that logistic-style replication dynamics (Eq. (12)) still yields growth that approximately exponential. As shown in Fig. 1, however, this near-exponential growth does not match the pronounced non-exponential growth modes (linear, super-exponential, or convex) observed in several bacterial species. Additional time-dependent transcriptional regulation is therefore necessary to reproduce those distinctly non-exponential growth phenotypes.

## V. MECHANISTIC ORIGINS OF NON-EXPONENTIAL GROWTH

Gene expression can vary extensively due to environmental changes [31, 36] and transiently during gene duplication [42, 43]. Since simple proportionality between transcription rate and DNA content fails to reproduce experimentally observed dynamics of elongation rate *κ*_*L*_ (compare Fig. 1 and Fig. 3), we next consider how deviations from DNA-proportional transcription (Eq. 13) affect growth rate dynamics. Specifically, we explore cell-cycle periodic perturbations of the transcription rates *ω*_*q*_(*t*) that preserve steady-state growth across generations. Physically, these perturbations could arise from changes in RNA polymerase concentration or promoter activity. Although our model does not explicitly track gene regulators, it captures how altered probabilities of transcriptional “bursting” events translate to changes in average growth rate.

In Section V A, we illustrate the qualitative impact of gene-expression perturbations using a square-wave perturbation to the transcription rate. Next, Section V B investigates how sinusoidal variations in ribosome and envelope transcription at different times and amplitudes within the cell cycle impact growth rate. We then examine, in Section V C, the timescales on which these regulatory changes influence growth. Finally, Section VI uses these insights to elucidate the diverse non-exponential growth patterns observed in different bacterial species.

### A. Deviation from DNA-proportional Transcription

To understand how dynamic changes in transcription rate affect growth dynamics, we first consider a simple, square-wave perturbation to the transcription rate with clearly defined “on” and “off” phases. Specifically, let

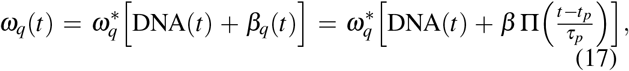

where *β*_*q*_(*t*) is the perturbation term, Π is the Heaviside (step) function, *τ*_*p*_ the pulse duration, and *t*_*p*_ the pulse initiation time. Thus, at time *t*_*p*_, transcription rate of sector *q* increases by an amount *β* for a period of length *τ*_*p*_, after which it returns to the baseline 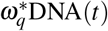.Figures 4a,b illustrate this perturbation for the ribosome sector (*β*_*R*_ *≠*0, *β*_*E*_ = 0), and Figs. 4c,d do so for the envelope sector (*β*_*R*_ ≠ 0, *β*_*E*_ = 0).

**FIG. 4.**
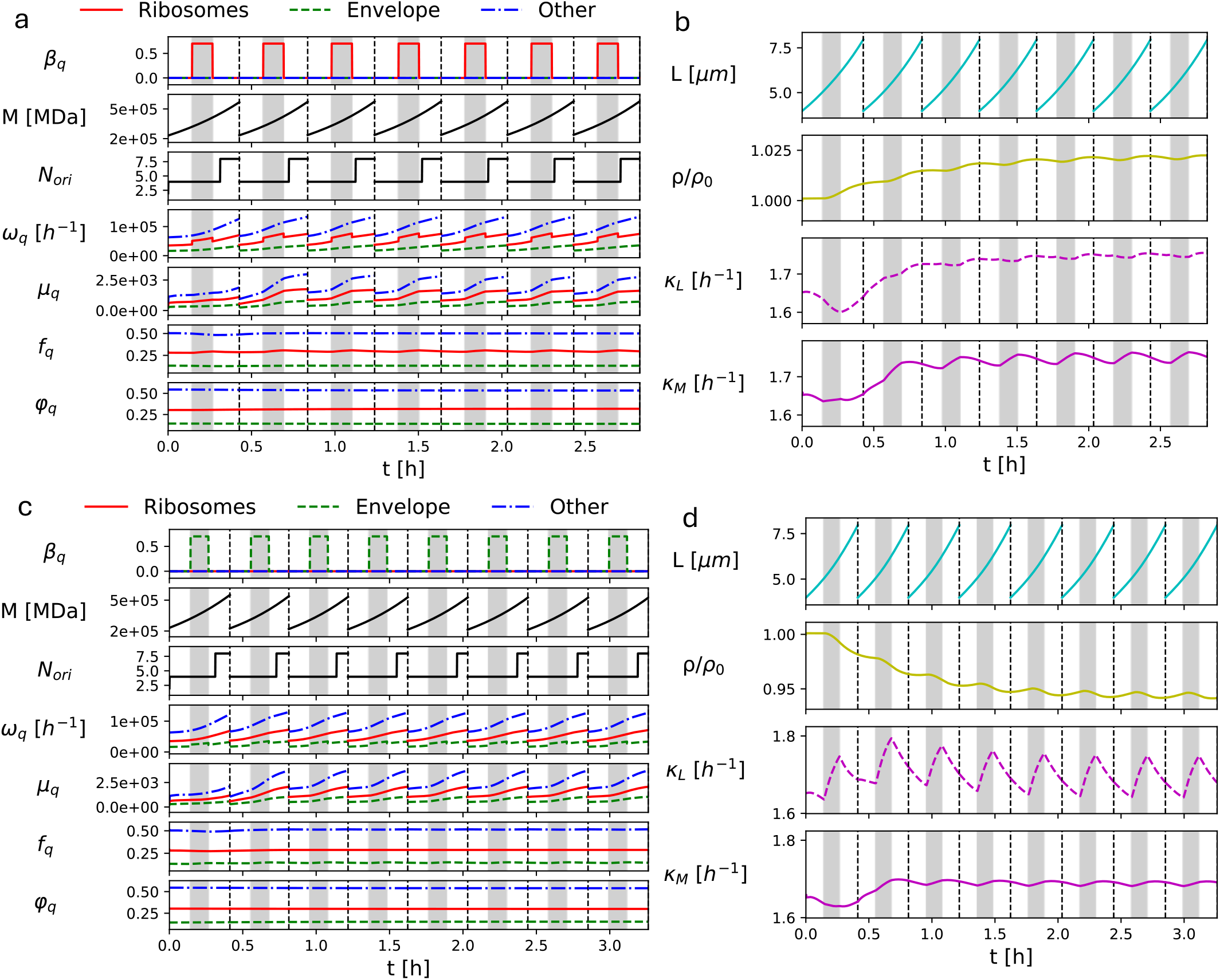
Effects of transcription perturbations on cell growth. (a) Trajectories of physiological variables evolved in the model (top to bottom): perturbation to transcription rates *β*_*q*_, cell mass *M*, the number of replication origins *N*_*ori*_, transcription rates *ω*_*q*_, mRNA abundance *µ*_*q*_, proteomme allocation fraction *f*_*q*_ and mass fraction *φ*_*q*_. Here we only consider a square wave perturbation to the transcription rate of ribosomal proteins in absolute time. The square wave perturbation is given by Eq. (17) with the parameters *t*_*p*_ = 0.5*τ, τ*_*p*_ = 0.3*τ*, and *β* = 0.7. This period is illustrated with a grey region in each cell cycle. (b) Model output showing the trajectory of (top to bottom) cell length *L*, normalized cell density *ρ/ρ*_0_, elongation rate *κ*_*L*_, and mass growth rate *κ*_*M*_ for the simulation corresponding to data in panel (a). Density is relative to the initialized value *ρ* = 1.9 · 10^5^ MDa*/µ*m^3^ [50]. Elongation and growth rates are calculated according to Eq. (15) and Eq. (16), respectively. (c) and (d) show an identical perturbation, but to the expression of cell envelope proteins *ω*_*E*_ rather than the cell envelope.

In the case of perturbation to the transcription rate of ribosomal proteins (Figs. 4a,b), *f*_*R*_ rises during the pulse and returns to its initial level afterward. Consequently, the mass growth rate *κ*_*M*_ increases while the elongation rate *κ*_*L*_ decreases, reflecting the trade-off between ribosomal (*f*_*R*_) and envelope (*f*_*E*_) proteome allocation. The cell density temporarily increases during the perturbation and remains slightly elevated on average compared to the unperturbed steady state due to the periodic nature of the perturbation. Notably, changing *ω*_*R*_ affects *κ*_*M*_ more strongly than *κ*_*L*_, emphasizing the ribosomal sector’s primary role in controlling mass growth rate.

By contrast, perturbations to the transcription of envelope proteins (Figs. 4c,d) increases *f*_*E*_ with minimal impact on *f*_*R*_, leading to an increase in *κ*_*L*_. Although the initial cycle shows a mild decrease in *κ*_*M*_, this effect is partially offset by the adder-like division criterion on *M*_*E*_, which triggers division at a lower total mass and lowers the overall cell density in subsequent cell cycles. Consequently, perturbing *ω*_*E*_ predominantly influences *κ*_*L*_ rather than *κ*_*M*_, underscoring how envelope allocation principally drives cell elongation. In both scenarios, resource allocation constraints introduce secondary feedbacks, but the primary effect remains sector-specific: perturbing ribosome transcription chiefly alters bulk mass growth, whereas envelope transcription primarily affects elongation.

### B. Cyclical Gene Expression

Having established a basic intuition for how transcriptional fluctuations shape growth, we now introduce a biologically motivated form of time-dependence of transcription rate. Since gene expression must be cell-cycle periodic at steady state, many functions could, in principle, model such oscillatory behavior. To compare scenarios where transcription changes primarily in the first versus the second half of the cycle, we use a simple sinusoidal perturbation:

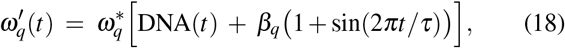

where *β*_*q*_ sets the perturbation amplitude for sector *q*. Such periodic regulation may arise from diverse biological mechanisms, including promoter-specific regulation or feedback loops tied to replication, division, and morphogenesis [25, 34]. Unlike the square-wave peerturbation, this smooth, continuous function better approximates the average outcome of noisy biochemical processes.

Figure 5a shows an example perturbation of this form, where *ω*_*R*_ is increased in the first half of the cell cycle, while *ω*_*E*_ is increased in the second half. Correspondingly, Fig. 5b shows that the elongation rate *κ*_*L*_ decreases and the mass growth rate *κ*_*M*_ increases during the interval of higher ribosome expression, and vice versa for envelope expression. This leads to a local maximum in *κ*_*M*_ roughly at mid-cycle, shortly after a minimum in *κ*_*L*_.

**FIG. 5.**
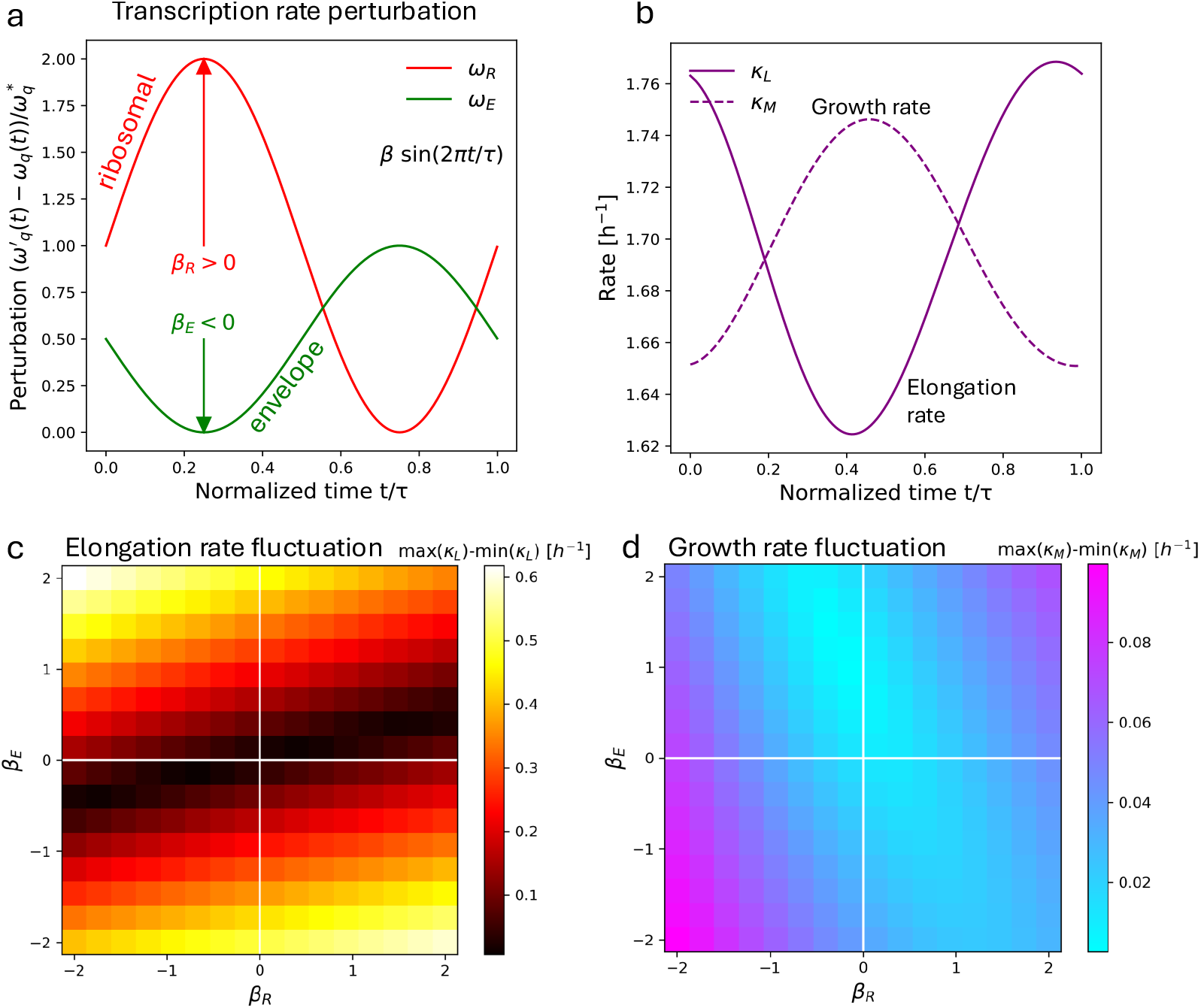
Effects of cyclical gene expression on growth. (a) A representative set of sinusoidal perturbations to *ω*_*R*_(*t*) and *ω*_*E*_ (*t*) as defined in Eq. (18) plotted against normalized time *t/τ*, where *τ* is cell cycle duration. Note that the sign of *β* determines whether or not the phase is offset by *π*. (b) Elongation and growth rates corresponding to the perturbation in (a) applied to the cell simulated in Fig. 3: *κ*_*n*,0_ = 25h^−1^, *ω*_*R*_ = 1.77 ·10^4^h^−1^, *ω*_*E*_ = 8.45 ·10^3^h^−1^, and *ω*_0_ = 3.18 ·10^4^h^−1^ with a steady-state tolerance of 1%. (c) A colormap showing the range of the elongation rate during the cell cycle. The y-axis corresponds to the magnitude (|*β*|) and phase (sign of *β*) of the envelope perturbation and the x-axis corresponds to the ribosomal transcription perturbation. The white lines denote the quadrants in which the perturbations are either in or out of phase with each other. Cell-cycle averaged mass fractions and cell cycle time are constrained to be the same as the unperturbed state for each entry by fitting *ω*_*R*_, *ω*_*E*_, and *ω*_0_. (d) A colormap showing the range of the mass growth rate during the cell cycle with the same axes as (c).

To explore these effects systematically, we map how *β*_*R*_ and *β*_*E*_ influence the range of *κ*_*L*_ and *κ*_*M*_ over each cell cycle (Figs. 5c,d). As with the square-wave perturbations, changes in *ω*_*R*_ primarily affect *κ*_*M*_, while shifts in *ω*_*E*_ principally alter *κ*_*L*_. However, their relative phases also matter. When ribosome and envelope perturbations are out-of-phase (opposite signs for *β*_*R*_ and *β*_*E*_), the elongation rate exhibits the largest changes, owing to trade-offs in allocation. By contrast, inphase perturbations (same signs for *β*_*R*_ and *β*_*E*_) yield more pronounced variations in *κ*_*M*_ as such perturbations result in oscillations of the total transcription rate. Notably, these fluctuations differ by an order of magnitude, with *κ*_*L*_ fluctuations typically on the order of 10^−1^ h^−1^ and *κ*_*M*_ fluctuations on the order of 10^−2^ h^−1^. Likewise, variations in *κ*_*L*_ from changing *ω*_*E*_ generally exceed those from altering *ω*_*R*_. Thus, the timing and amplitude of gene-expression oscillations can systematically reshape both mass and length growth rates.

### C. Timescale of Gene-Expression Perturbations and Growth-Rate Response

Although time-dependent changes in gene expression lead to deviations from exponential growth, such transcriptional perturbations do not instantaneously alter growth rates. Instead, they must propagate through multiple molecular layers before manifesting in observable changes to mass or elongation rates. We thus elucidate how quickly (or slowly) these perturbations translate into changes in elongation (*κ*_*L*_) and mass growth (*κ*_*M*_) rates. Our simulations reveal stark contrasts: transcriptional changes typically take hours to manifest in our coarse-grained sectors, whereas proteome allocation shifts can elicit immediate responses.

To illustrate this, consider a step-function decrease in *ω*_*R*_ (Fig. 6a). Even though ribosomal transcription declines abruptly, the effect on *κ*_*M*_ unfolds over several hours, as the altered ribosome production must propagate through both the transcriptome and proteome. Meanwhile, *κ*_*L*_ temporarily increases owing to a relative boost in envelope expression, but it too settles to the new steady state only after around four hours. A similar step decrease in *ω*_*E*_ (Fig. 6b) triggers an immediate reduction in *κ*_*M*_, while *κ*_*L*_ first falls to a minimum, then ramps back up over a comparable timescale.

**FIG. 6.**
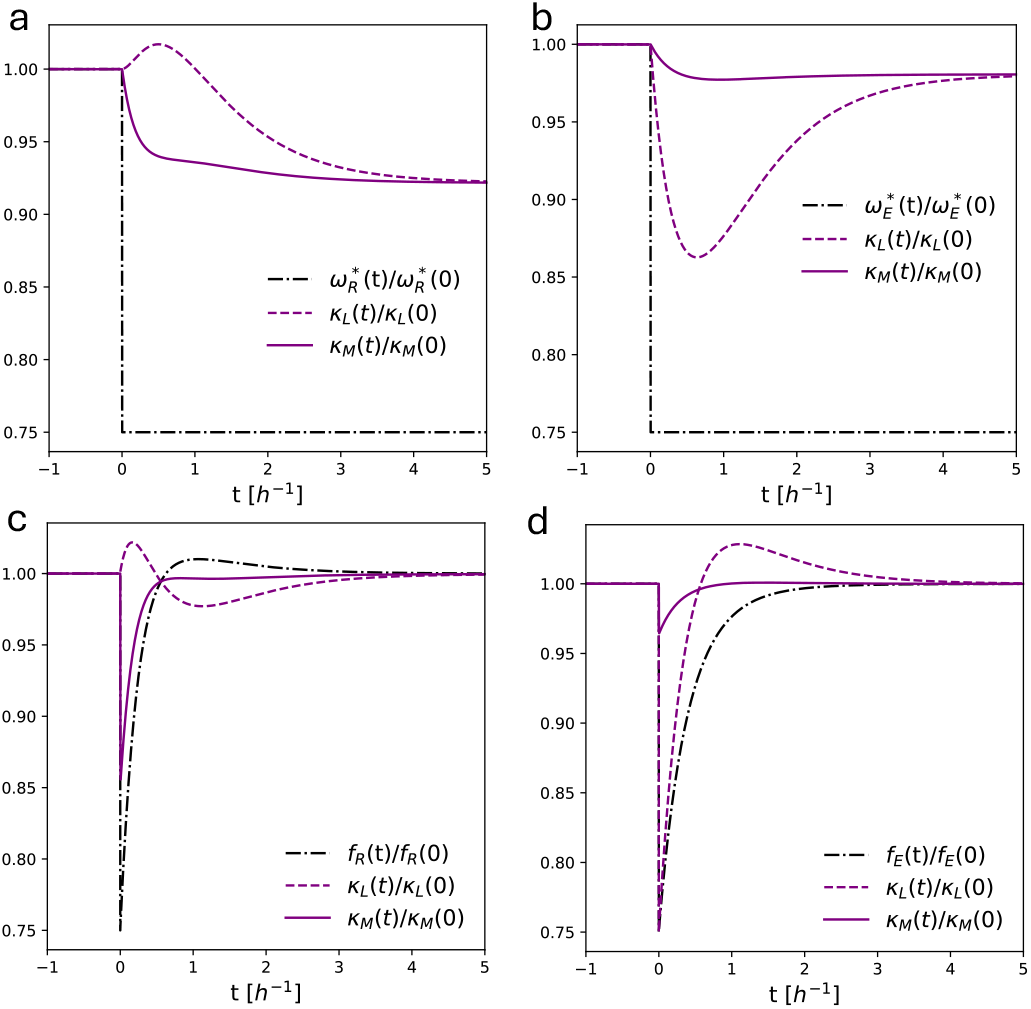
Timescale limitations of the effects of gene expression on growth. (a) A simulation of balanced exponential growth with *ω*(*t*) determined by Eq. (13). At 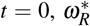 is decreased to 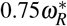.We plot the effects of this transcription perturbation on elongation and growth rates, as calculated by Eq. (15) and Eq. (16). (b) Considers an identical scenario to (a) except with a step decrease in 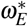.(c) Simulating the effects of a step decrease in *f*_*R*_ by a factor of 0.75, on cell growth and elongation rates. Note that *f*_*R*_ relaxes back to the initial value as *ω*_*R*_ remains unchanged. (d) Shows the effects of a step decrease in *f*_*E*_.

In contrast, altering proteome allocation fractions *f*_*q*_ produces rapid shifts in growth rates (Figs. 6c,d). A sudden drop in *f*_*R*_ (with *f*_*F*_ rising accordingly) instantly lowers *κ*_*M*_ and boosts *κ*_*L*_, since a larger fraction of ribosomes is devoted to envelope synthesis. This transient peak in *κ*_*L*_ emerges in half the time required by the transcriptional perturbation, and the system then returns to the original steady state. Likewise, reducing *f*_*E*_ prompts an immediate decrease in both *κ*_*L*_ and *κ*_*M*_, followed by an overshoot and a return to equilibrium within roughly half an hour.

Thus transcriptional regulation operates on hours-long timescales, reflecting the need for new mRNA transcripts and protein turnover. By contrast, direct changes to proteome allocation fractions can rewire growth rates almost instantaneously. This disparity underscores the multi-layered nature of bacterial growth control, wherein slower upstream processes (transcription) and faster downstream processes (allocation) together shape the overall dynamics of cell growth.

## VI. MODELING NON-EXPONENTIAL ELONGATION MODES IN BACTERIAL CELLS

### A. Convex growth of *B. subtilis*

The periodic transcription perturbations described in Section V B provide a framework for understanding the convex growth pattern of *Bacillus subtilis*. Experimentally, the elongation rate of *B. subtilis* has been observed to change by 0.18 h^−1^ to 0.52 h^−1^ across the cell cycle [8]. From Fig. 5, we see that achieving such a substantial temporal variation in *κ*_*L*_ requires a pronounced, cell-cycle-dependent increase in envelope transcription *ω*_*E*_. Whether or not ribosome transcription *ω*_*R*_ must also vary remains to be seen; however, the data strongly suggest that growth-related envelope genes do not remain at a fixed transcription level throughout the cycle.

To quantitatively understand the behavior of *ω*_*R*_, we fit a sinusoidal perturbation (generalized from Eq. (18) to include a phase shift) to the ensemble-averaged elongation rate data in Fig. 1b:

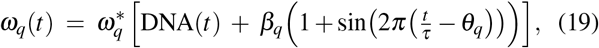

where the parameter *θ*_*q*_ sets the phase. Figure 7a shows an example fit of this form. We see that, as expected, there is an increase in *ω*_*E*_ during the later half of the cell cycle when *κ*_*L*_ increases in the data (Fig. 7b,c). The fit also indicates a corresponding trade-off in *ω*_*R*_; rather than adjusting *ω*_*E*_ alone, ribosome transcription is front-loaded in the cycle while envelope transcription is back-loaded. As seen previously in Fig. 5c, these out-of-phase perturbations enhance changes in *κ*_*L*_ without altering the cell’s average mass fractions or cycle time.

**FIG. 7.**
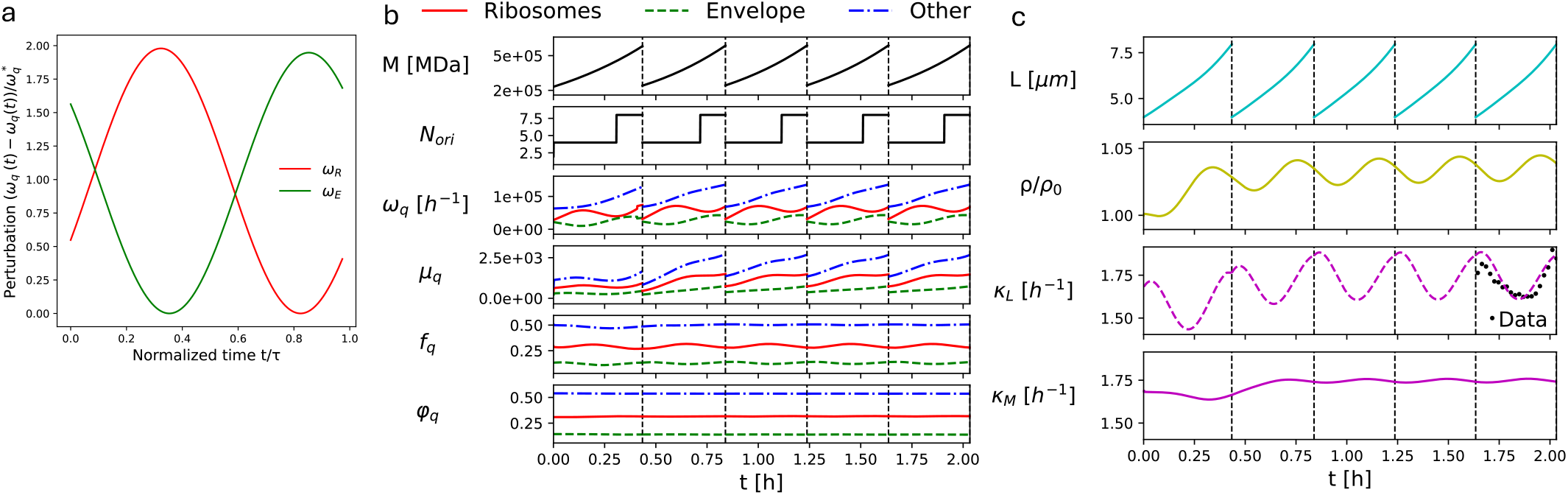
Modeling convex elongation rate profile of *B. subtilis*. (a) Perturbations to ribosome and cell envelope transcriotion rates, of the form in Eq. (19), plotted against normalized time *t/τ*, where *τ* is cell cycle duration. The perturbation profile is fited to the elongation rate data of *B. subtilis* cells (see panel (c)). Fitting parameters are *β*_*R*_ = 0.99, *β*_*E*_ = 0.97, *θ*_*R*_ = 0.93h, and *θ*_*E*_ = 0.40h. (b) Trajectories of the cell physiological variables predicted by model simulation, corresponding to the perturbations shown in panel (a). The parameters used are *κ*_*n*,0_ = 25h^−1^, *ω*_*R*_ = 1.77 · 10^4^h^−1^, *ω*_*E*_ = 8.45 · 10^3^h^−1^, and *ω*_0_ = 3.18 · 10^4^h^−1^. (c) Simulation outputs for cell length, density, elongation rate and growth rate for the simulation results presented in (b). Density is relative to the initialized value *ρ* = 1.9 10 MDa*/µ*m [50]. Elongation and growth rates are calculated according to Eq. (15) and Eq. (16), respectively. Elongation rate data are for *B. subtilis* BSB1 cells grown in Glycerol at 37°*C* [8].

Interestingly, despite both *ω*_*R*_ and *ω*_*E*_ being perturbed, *κ*_*M*_ remains nearly constant compared to the pronounced mid-cycle dip and rise in *κ*_*L*_. This implies that the experimentally observed convex elongation rate profile does not reflect dramatic variations in the overall mass growth rate. Instead, the envelope-specific transcription dynamics cause minor cyclic fluctuations in the cell’s dry mass density, producing a convex elongation pattern in real time. This distinction underscores how subtle variations in sector-specific gene expression can manifest as noticeable cell-shape changes, even when total biomass growth remains close to exponential.

### B. Super- and sub-exponential growth

Both super-exponential and linear growth exhibit a jump discontinuity at division (Fig. 1a) as, on average, the elongation rate resets to its value at the start of the previous cell cycle. In Section V C, we showed that an instantaneous change in gene expression at division cannot, by itself, generate this instantaneous shift in elongation rate on the short (~ 1 minute) timescale. Given that cell division is symmetric on average for *E. coli* [4], the only remaining mechanism is a change in proteome allocation fractions due to division itself.

Focusing on the monotonic changes in elongation rate observed in super-exponential and linear growth, suppose that the simplest possible time dependence in proteome allocation fractions is a linear increase (or decrease) throughout the cell cycle, followed by a jump discontinuity at division. In that case, we can write

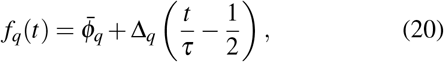

where 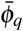 is the cycle-averaged mass fraction, *τ* is the cell cycle time, and Δ_*q*_ is the overall change in *f*_*q*_ over one cycle. Fitting Eq. (20) to *E. coli* data [4] (Fig. 8a,b) shows that *f*_*E*_ grows throughout the cell cycle and then returns to its initial value at the next generation, mirroring a previous super-exponential growth model [27]. By contrast, fitting the same form to *M. tuberculosis* data [42] implies a decrease in envelope allocation (*f*_*E*_) over the cell cycle—indicating that super-exponential growth arises from a relative increase in allocation to ribosomes, whereas linear growth is driven by a relative decrease.

**FIG. 8.**
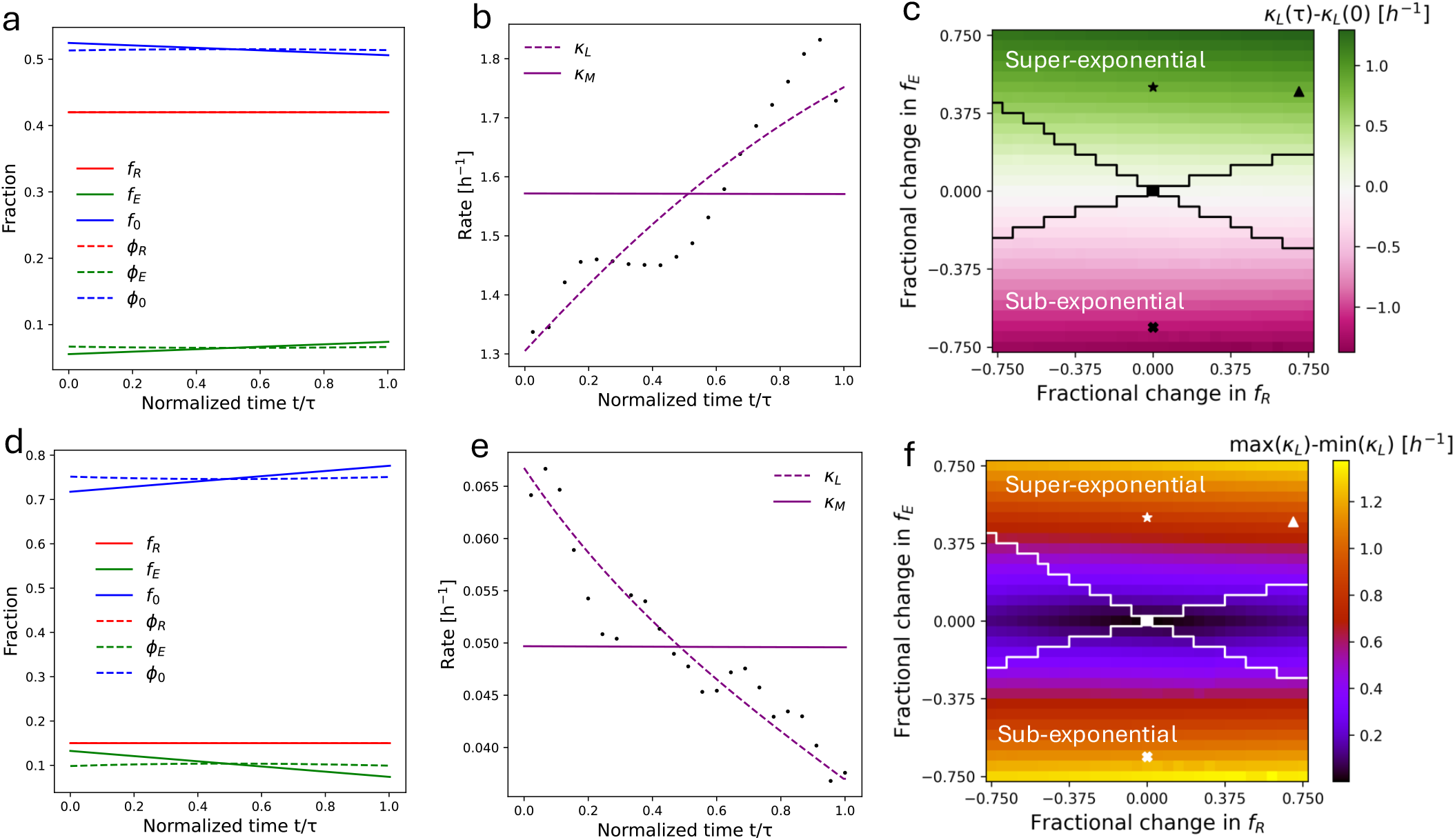
Dynamic resource allocation during super- and sub-exponential elongation. (a) Ribosome allocation fractions (solid lines, Eq. (20)) and integrated mass fractions (dashed lines) vs. normalized time *t/τ*, fitted to *E. coli* super-exponential growth rate data [4]. The fitted parameter values are: 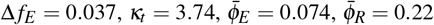 [31]. (b) Elongation (Eq. (15)) and mass growth rates (Eq. (16)) corresponding to the allocation in (a), plotted against normalized time *t/τ*. The plotted points show the ensemble-averaged growth rate data for *E. coli* K12 NCM3722 cells, grown in supplemented Glucose media at 37°*C* [4]. (c) Heatmap of the net change in elongation rate from birth (*t* = 0) to division (*t* = *τ*), varying the fractional changes in envelope (Δ *f*_*E*_) and ribosome (Δ *f*_*R*_) allocation. The diagonal line divides super-exponential (top) and sub-exponential (bottom) regimes; left and right regions denote monotonic vs. non-monotonic growth. The star marks the *E. coli* fit in (a,b), the triangle marks a variant with 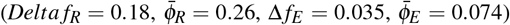.the cross marks the linear fit in (d,e), and the black square at the origin denotes purely exponential growth. (d) Analogous resource allocation fit for *M. tuberculosis* 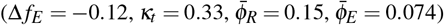. (e) Corresponding elongation and mass growth rates, plotted with *M. tuberculosis* data [30]. (f) Heatmap of the total range of elongation rate over one cycle, labeled as in (c).

Figure 8c illustrates how the change in *κ*_*L*_ from birth to division depends on Δ_*R*_ and Δ_*E*_. Positive Δ_*E*_ leads to super-exponential growth, negative Δ_*E*_ to sub-exponential growth (with linear growth as a special case). Once Δ_*E*_ is fixed, Δ_*R*_ has minimal effect on the overall change in *κ*_*L*_, meaning that changes in *f*_*E*_ primarily drive growth on the timescale of a single cell cycle. However, Δ_*R*_ does influence the concavity of the *κ*_*L*_ trajectory, yielding slight non-monotonic regions if Δ_*R*_ is large compared to Δ_*E*_. Figure 8f shows that these non-monotonic deviations remain small and do not produce full oscillations, agreeing with our discussion in Section V B. Thus, Δ_*E*_ sets the amplitude of the *κ*_*L*_ variation, while Δ_*R*_ determines whether the curve is concave or convex. When Δ_*R*_ *>* 0, the trajectory is convex, in contrast to the concave shape shown in Fig. 8b for Δ_*R*_ ≤ 0. Since super-exponential growth in filamentous *E. coli* is known to approach exponential growth asymptotically [27] (i.e., a concave shape), our model supports the scenario of Δ_*R*_ ≤ 0. In short, we predict a trade-off in proteome allocation: ribosome synthesis is emphasized early in the cycle, while envelope synthesis is prioritized later.

Returning to the full gene expression-driven model, Fig. 6c,d shows that a step-like change to *f*_*R*_ or *f*_*E*_ (e.g., at division) without transcriptional adjustments simply relaxes back to steady state on the same timescale that filamentous cells reach steady-state growth [27]. This indicates that time-dependent gene expression is not strictly necessary to produce super-exponential growth; a division-based reallocation alone can suffice.

## VII. DISCUSSION

Recent single-cell measurements reveal that bacterial elongation diverges from the standard assumption of exponential growth, instead showing convex, super-exponential, or sub-exponential elongation rate profiles across species. To explain these patterns, we built a minimal mechanistic model linking transcription and proteome allocation to to overall mass growth and elongation rate. Under the simplest assumption of mRNA synthesis rate proportional to gene dosage, the model predicts near-exponential growth in cell size and mass —which fails to capture the observed modes of elongation rates in single-cell data. Consequently, we explored how deviations from DNA-proportional transcription could shape growth dynamics.

In particular, we introduced cell-cycle periodic perturbations to transcription and analyzed their effects. Altering ribosome expression strongly influences mass growth rate, while envelope expression modulations most profoundly affect elongation rate. When total transcription shifts in-phase, mass growth changes significantly; however, out-of-phase peaks in ribosome and envelope expression lead to more dramatic fluctuations in elongation. Crucially, envelope-specific perturbations shift elongation by roughly an order of magnitude more than ribosomal changes do. Fitting our model to *B. subtilis* data shows that convex elongation rate requires envelope allocation to change over the cell cycle, balanced by a trade-off with ribosome synthesis.

We next turned to how some species achieve super-exponential or sub-exponential growth with apparent jump discontinuities at division. Our analysis shows that instantaneous changes in gene expression alone cannot produce such sharp transitions. Instead, a division-induced shift in resource allocation is required, resetting envelope synthesis at each division event. Super-exponential growth follows when envelope allocation increases throughout the cycle but dips at division, whereas sub-exponential growth occurs when envelope allocation decreases over the cycle and then rebounds at division. This mechanism aligns with observations in *E. coli*, where non-growing poles reduce the actively elongating surface area immediately post-division, which subsequently expands as cells elongate. In contrast, *M. tuberculosis*—which elongates only from its poles—shows a surge in envelope allocation upon activating new poles after division, then tapers as the cell elongates.

While our model captures the core aspects of cell-cycle-dependent growth rate profile, it does not yield a unique transcription-time profile from elongation data alone. Future experiments quantifying single-cell transcription, dry mass density, and translational activity (e.g., via ribosome profiling) could refine these predictions. Additionally, we employ a minimal proteome perspective, omitting detailed functions of complexes such as the elongasome or divisome and ignoring potential noise in transcription or division timing. Still, our findings highlight how non-exponential elongation can emerge from coordinated changes in envelope and ribosome expression, coupled with division-driven resource reallocation. Ultimately, the evolutionary or fitness benefits of non-exponential elongation remain unclear, since deviations from balanced, exponential growth appear to exert only minor effects on cell cycle timing. Nonetheless, understanding why cells deviate from perfectly exponential growth, even when balanced biosynthesis seems optimal, can reveal intragenerational physiological strategies that remain hidden in population-averaged data.

## ACKNOWLEDGMENT

This work is supported by the National Institutes of Health (NIH R35 GM143042) and the Shurl and Kay Curci Foundation.

## References

[1] J. Monod, The growth of bacterial cultures, Annual Review of Microbiology 3, 371 (1949).

[2] P. Wang, L. Robert, J. Pelletier, W. L. Dang, F. Taddei, A. Wright, and S. Jun, Robust growth of escherichia coli, Current Biology 20, 1099 (2010).

[3] Y. Lin, S. Crosson, and N. F. Scherer, Single-gene tuning of caulobacter cell cycle period and noise, swarming motility, and surface adhesion, Molecular Systems Biology 6, 445 (2010).

[4] S. Taheri-Araghi, S. Bradde, J. T. Sauls, N. S. Hill, P. A. Levin, J. Paulsson, M. Vergassola, and S. Jun, Cell-size control and homeostasis in bacteria, Current Biology 25, 385 (2015).

[5] C. S. Wright, S. Banerjee, S. Iyer-Biswas, S. Crosson, A. R. Dinner, and N. F. Scherer, Intergenerational continuity of cell shape dynamics in caulobacter crescentus, Scientific Reports 5, 1 (2015).

[6] Y. Tanouchi, A. Pai, H. Park, S. Huang, N. E. Buchler, and L. You, Long-term growth data of escherichia coli at a single-cell level, Scientific Data 4, 1 (2017).

[7] L. Susman, M. Kohram, H. Vashistha, J. T. Nechleba, H. Salman, and N. Brenner, Individuality and slow dynamics in bacterial growth homeostasis, Proceedings of the National Academy of Sciences 115, E5679 (2018).

[8] J. T. Sauls, S. E. Cox, Q. Do, V. Castillo, Z. Ghulam-Jelani, and S. Jun, Control of bacillus subtilis replication initiation during physiological transitions and perturbations, mBio 10, 10.1128/mbio.02205-19 (2019).

[9] A. Amir, Cell size regulation in bacteria, Physical Review Letters 112, 208102 (2014).

[10] M. Campos, I. Surovtsev, S. Kato, A. Paintdakhi, B. Beltran, S. Ebmeier, and C. Jacobs-Wagner, A constant size extension drives bacterial cell size homeostasis, Cell 159, 1433–1446 (2014).

[11] S. Jun and S. Taheri-Araghi, Cell-size maintenance: universal strategy revealed, Trends in Microbiology 23, 4 (2015).

[12] S. Banerjee, K. Lo, M. K. Daddysman, A. Selewa, T. Kuntz, A. R. Dinner, and N. F. Scherer, Biphasic growth dynamics control cell division in caulobacter crescentus, Nature Microbiology 2, 1 (2017).

[13] S. Jun, F. Si, R. Pugatch, and M. Scott, Fundamental principles in bacterial physiology—history, recent progress, and the future with focus on cell size control: a review, Reports on Progress in Physics 81, 056601 (2018).

[14] F. Si, G. Le Treut, J. T. Sauls, S. Vadia, P. A. Levin, and S. Jun, Mechanistic origin of cell-size control and homeostasis in bacteria, Current Biology 29, 1760 (2019).

[15] F. Si, D. Li, S. E. Cox, J. T. Sauls, O. Azizi, C. Sou, A. B. Schwartz, M. J. Erickstad, Y. Jun, X. Li, et al., Invariance of initiation mass and predictability of cell size in escherichia coli, Current Biology 27, 1278 (2017).

[16] H. Zheng, Y. Bai, M. Jiang, T. A. Tokuyasu, X. Huang, F. Zhong, Y. Wu, X. Fu, N. Kleckner, T. Hwa, et al., General quantitative relations linking cell growth and the cell cycle in escherichia coli, Nature Microbiology 5, 995 (2020).

[17] D. Serbanescu, N. Ojkic, and S. Banerjee, Cellular resource allocation strategies for cell size and shape control in bacteria, The FEBS Journal 289, 7891 (2022).

[18] L. K. Harris and J. A. Theriot, Relative rates of surface and volume synthesis set bacterial cell size, Cell 165, 1479 (2016).

[19] N. Ojkic, D. Serbanescu, and S. Banerjee, Surface-to-volume scaling and aspect ratio preservation in rod-shaped bacteria, eLife 8, e47033 (2019).

[20] E. R. Oldewurtel, Y. Kitahara, and S. van Teeffelen, Robust surface-to-mass coupling and turgor-dependent cell width determine bacterial dry-mass density, PNAS 10.1073/pnas.2021416118 (2021).

[21] S. Banerjee, K. Lo, N. Ojkic, R. Stephens, N. F. Scherer, and A. R. Dinner, Mechanical feedback promotes bacterial adaptation to antibiotics, Nature Physics 17, 403 (2021).

[22] J. Nguyen, V. Fernandez, S. Pontrelli, U. Sauer, M. Ackermann, and R. Stocker, A distinct growth physiology enhances bacterial growth under rapid nutrient fluctuations, Nature Communications 12, 1 (2021).

[23] M. Panlilio, J. Grilli, G. Tallarico, I. Iuliani, B. Sclavi, P. Cicuta, and M. Cosentino Lagomarsino, Threshold accumulation of a constitutive protein explains e. coli cell-division behavior in nutrient upshifts, Proceedings of the National Academy of Sciences 118, e2016391118 (2021).

[24] S. Bakshi, E. Leoncini, C. Baker, S. J. Cañas-Duarte, B. Okumus, and J. Paulsson, Tracking bacterial lineages in complex and dynamic environments with applications for growth control and persistence, Nature Microbiology 6, 783 (2021).

[25] J. Kratz and S. Banerjee, Dynamic proteome trade-offs regulate bacterial cell size and growth in fluctuating nutrient environments, Communications Biology 6 (2023).

[26] P. Kar, S. Tiruvadi-Krishnan, J. Männik, J. Männik, and A. Amir, Distinguishing different modes of growth using single-cell data, eLife 10, e72565 (2021).

[27] A. Cylke and S. Banerjee, Super-exponential growth and stochastic size dynamics in rod-like bacteria, Biophysical Journal 122, 1254–1267 (2023).

[28] T. W. Ng, N. Ojkic, D. Serbanescu, and S. Banerjee, Differential growth regulates asymmetric size partitioning in caulobacter crescentus, Life Science Alliance 7, 10.26508/lsa.202402591 (2024).

[29] N. Nordholt, J. H. van Heerden, and F. J. Bruggeman, Biphasic cell-size and growth-rate homeostasis by single bacillus subtilis cells, Current Biology 30, 2238 (2020).

[30] E. S. Chung, P. Kar, M. Kamkaew, A. Amir, and B. B. Aldridge, Single-cell imaging of the mycobacterium tuberculosis cell cycle reveals linear and heterogenous growth, Nature Microbiology 9, 3332–3344 (2024).

[31] M. Scott, C. W. Gunderson, E. M. Mateescu, Z. Zhang, and T. Hwa, Interdependence of cell growth and gene expression: origins and consequences, Science 330, 1099 (2010).

[32] M. Scott, S. Klumpp, E. M. Mateescu, and T. Hwa, Emergence of robust growth laws from optimal regulation of ribosome synthesis, Molecular Systems Biology 10, 747 (2014).

[33] S. Hui, J. M. Silverman, S. S. Chen, D. W. Erickson, M. Basan, J. Wang, T. Hwa, and J. R. Williamson, Quantitative proteomic analysis reveals a simple strategy of global resource allocation in bacteria, Molecular Systems Biology 11, 784 (2015).

[34] A. Y. Weiße, D. A. Oyarzún, V. Danos, and P. S. Swain, Mechanistic links between cellular trade-offs, gene expression, and growth, Proceedings of the National Academy of Sciences 112, 10.1073/pnas.1416533112 (2015).

[35] N. Giordano, F. Mairet, J.-L. Gouzé, J. Geiselmann, and H. De Jong, Dynamical allocation of cellular resources as an optimal control problem: novel insights into microbial growth strategies, PLoS Computational Biology 12, e1004802 (2016).

[36] D. W. Erickson, S. J. Schink, V. Patsalo, J. R. Williamson, U. Gerland, and T. Hwa, A global resource allocation strategy governs growth transition kinetics of escherichia coli, Nature 551, 119 (2017).

[37] M. Mori, S. Schink, D. W. Erickson, U. Gerland, and T. Hwa, Quantifying the benefit of a proteome reserve in fluctuating environments, Nature Communications 8, 1 (2017).

[38] Y. K. Kohanim, D. Levi, G. Jona, B. D. Towbin, A. Bren, and U. Alon, A bacterial growth law out of steady state, Cell Reports 23, 2891 (2018).

[39] B. Alberts, R. Heald, A. Johnson, D. Morgan, M. Raff, K. Roberts, and P. Walter, Molecular biology of the cell, 5th ed. (W. W. Norton & Company, 2022).

[40] R. Balakrishnan, M. Mori, I. Segota, Z. Zhang, R. Aebersold, C. Ludwig, and T. Hwa, Principles of gene regulation quantitatively connect dna to rna and proteins in bacteria, Science 378, eabk2066 (2022),.

[41] J. Lin and A. Amir, Homeostasis of protein and mrna concentrations in growing cells, Nature Communications 9, 10.1038/s41467-018-06714-z (2018).

[42] A. W. Pountain, P. Jiang, T. Yao, E. Homaee, Y. Guan, K. J. McDonald, M. Podkowik, B. Shopsin, V. J. Torres, I. Golding, and et al., Transcription–replication interactions reveal bacterial genome regulation, Nature 626, 661–669 (2024).

[43] I. Golding and A. Amir, Gene expression in growing cells: A biophysical primer, arXiv 10.48550/arXiv.2311.12143 (2023).

[44] N. M. Belliveau, G. Chure, C. L. Hueschen, H. G. Garcia, J. Kondev, D. S. Fisher, J. A. Theriot, and R. Phillips, Fundamental limits on the rate of bacterial growth and their influence on proteomic composition, Cell Systems 12, 10.1016/j.cels.2021.06.002 (2021).

[45] A. Cylke, D. Serbanescu, and S. Banerjee, Energy allocation theory for bacterial growth control in and out of steady state, Proceedings of the Royal Society A: Mathematical, Physical and Engineering Sciences 480, 10.1098/rspa.2024.0219 (2024).

[46] Y. Taniguchi, Quantifying e. coli proteome and transcriptome with single-molecule sensitivity in single cells, Seibutsu Butsuri 51, 136–137 (2011).

[47] I. M. Keseler, Ecocyc: A comprehensive database resource for escherichia coli, Nucleic Acids Research 33, 10.1093/nar/gki108 (2004).

[48] D. N. Wilson and K. H. Nierhaus, The ribosome through the looking glass, Angewandte Chemie International Edition 42, 3464–3486 (2003).

[49] A. Tiessen, P. Pérez-Rodríguez, and L. J. Delaye-Arredondo, Mathematical modeling and comparison of protein size distribution in different plant, animal, fungal and microbial species reveals a negative correlation between protein size and protein number, thus providing insight into the evolution of proteomes, BMC Research Notes 5, 10.1186/1756-0500-5-85 (2012).

[50] Y. Kitahara, E. R. Oldewurtel, S. Wilson, Y. Sun, S. Altabe, D. de Mendoza, E. C. Garner, and S. van Teeffelen, The role of cell-envelope synthesis for envelope growth and cytoplasmic density in bacillus subtilis, PNAS Nexus 1, 10.1093/pnas-nexus/pgac134 (2022).

[51] A. Schmidt, K. Kochanowski, S. Vedelaar, E. Ahrné, B. Volkmer, L. Callipo, K. Knoops, M. Bauer, R. Aebersold, and M. Heinemann, The quantitative and condition-dependent escherichia coli proteome, Nature Biotechnology 34, 104 (2016).

